# Quantifying Seasonal and Diel Variation in Anopheline and Culex Human Biting Rates in Southern Ecuador

**DOI:** 10.1101/192773

**Authors:** Sadie J. Ryan, Catherine A. Lippi, Philipp H. Boersch-Supan, Naveed Heydari, Mercy Silva, Jefferson Adrian, Leonardo F. Noblecilla, Efraín B. Ayala, Mayling D. Encalada, David A. Larsen, Jesse T. Krisher, Lyndsay Krisher, Lauren Fregosi, Anna M. Stewart-Ibarra

## Abstract

**Background:** Quantifying mosquito biting rates for specific locations enables estimation of mosquito-borne disease risk, and can inform intervention efforts. Measuring biting itself is fraught with ethical concerns, so the landing rate of mosquitoes on humans is often used as a proxy measure. Southern coastal Ecuador was historically endemic for malaria (*P. falciparum* and *P. vivax*), although successful control efforts in the 2000s eliminated autochthonous transmission (since 2011). This study presents an analysis of data collected during the elimination period.

**Methods:** We examined human landing catch (HLC) data for three mosquito taxa: 2 malaria vectors, *Anopheles albimanus* and *Anopheles punctimacula*, and grouped *Culex spp*. These data were collected by the National Vector Control Service of the Ministry of Health over a 5-year time span (2007 – 2012) in five cities in southern coastal Ecuador, at multiple households, in all months of the year, during dusk-dawn (18:00-6:00) hours, often at both indoor and outdoor locations. Hurdle models were used to determine if biting activity was fundamentally different for the three taxa, and to identify spatial and temporal factors influencing bite rate. Due to the many different approaches to studying and quantifying bite rates in the literature, we also created a glossary of terms, to facilitate comparative studies in the future.

**Results:** Biting trends varied significantly with species and time. All taxa exhibited exophagic feeding behavior, and outdoor locations increased both the odds and incidence of bites across taxa. *An. albimanus* was most frequently observed biting, with an average of 4.7 bites per hour. The highest and lowest respective months for significant biting activity were March and July for *An. albimanus,* July and August for *An. punctimacula*, and February and July for *Culex spp.*

**Conclusions:** Fine-scale spatial and temporal differences exist in biting patterns among mosquito taxa in southern coastal Ecuador. This analysis provides detailed information for targeting vector control and household level behavioral interventions. These data were collected as part of routine vector surveillance conducted by the Ministry of Health, but such data have not been collected since. Reinstating such surveillance measures would provide important information to aid in preventing malaria re-emergence.

## Background

Despite major efforts to control and eliminate malaria and other vector-borne diseases through vector control, mosquito-borne diseases such as malaria, dengue, yellow fever, and now chikungunya and zika virus remain a major threat to people’s livelihoods in the Americas. An estimated 108 million people per year are at risk for malaria infections in the Americas, pointing to a need to maintain the eliminated status in areas that have successfully eradicated local infections and prevent reestablishment [1]. In Latin America there is high endemic diversity in both vectors and pathogens, including three species of malaria-causing parasites, *Plasmodium vivax*, *P. falciparium*, and *P. malariae* [1–4]. To monitor and measure the potential for mosquito-borne transmission, it is important to assess the risk or rate of infectious bites on humans. There are many challenges associated with the direct surveillance of pathogens, such as *Plasmodium*, in mosquito populations, thus vector-borne diseases are often monitored in terms of human case data [5–7]. The reliance on human cases to monitor vector-borne disease outbreaks is subject to many forms of reporting bias, and these biases may be further exacerbated in Ecuador, where disparities in clinical access may contribute to underreporting of cases, as is seen with dengue [8–10]. Even when clinical access is more widely available, as in urban areas, much of the public health data reported by Ecuador’s Ministry of Health relies on suspected clinical cases rather than laboratory confirmation [11]. Although malaria surveillance and diagnostics in Ecuador are much stronger relative to those of other mosquito-borne diseases, detection of asymptomatic malaria and cases in remission remain a challenge to surveillance and disease elimination [12,13].

Measuring force of infection, or transmission risk of mosquito-borne diseases through models of vital rates [14–17], require knowledge of many components of the transmission cycle, including biting rates. The Entomological Inoculation Rate (EIR) is commonly used as a means of describing potential risk of infection from vectorborne diseases; this is the rate of infectious bites per person per day, usually estimated, or derived from biting rates and a measure of vector infection prevalence. EIR is considered a more direct measure of infection intensity than human incidence or other traditional epidemiological measures [18,19]. Clearly, measuring the rate of infection in vectors can be logistically complex, but capturing an estimate of biting rate, perhaps less so. Thus, a simplified attempt to quantify potential disease transmission is the development of human bite rate (HBR) and landing rate (LR) indices, generally described as the number of mosquitoes of a species respectively exhibiting feeding or resting behavior on a human recorded for a given location and time period [20–22]. Although most commonly used in the context of establishing the number of female mosquitoes that are attempting to take blood meals in field or laboratory conditions, there is a great deal of variability in the literature with regards to the definitions and field protocols associated with these metrics.

With this in mind, we developed a glossary of terms we encountered in the literature, to facilitate communication of definitions, as a means to both measure and interpret study findings for comparison (Table 1). In general, the protocol for HBR and LR studies involves an initial survey for potential sites, a species inventory to establish vector presence, training field entomology technicians in identification of species and behaviors, and establishing spatial points and temporal intervals for data collection [23]. Like raw mosquito density, HBR and LR do not directly measure infections, but these indices are often cited as a proxy for species presence, density of blood-seeking females, and the capacity for disease transmission [23,24]. Potential issues with HBR include reliance on visual identification of mosquito species, inter-observer agreement, and exposure of workers to pathogens [25–28]. Human landing catch (HLC), wherein mosquitoes counted in the landing rate survey are captured and later examined in the lab, can overcome most of these obstacles, but at the cost of additional field and laboratory resources [22]. Depending on study design and data collection protocol, bite rate indices have the potential to provide a wealth of information regarding vector behavior at very fine spatial and temporal scales in a manner that is both relatively cost-effective and efficient.

**Table 1.**
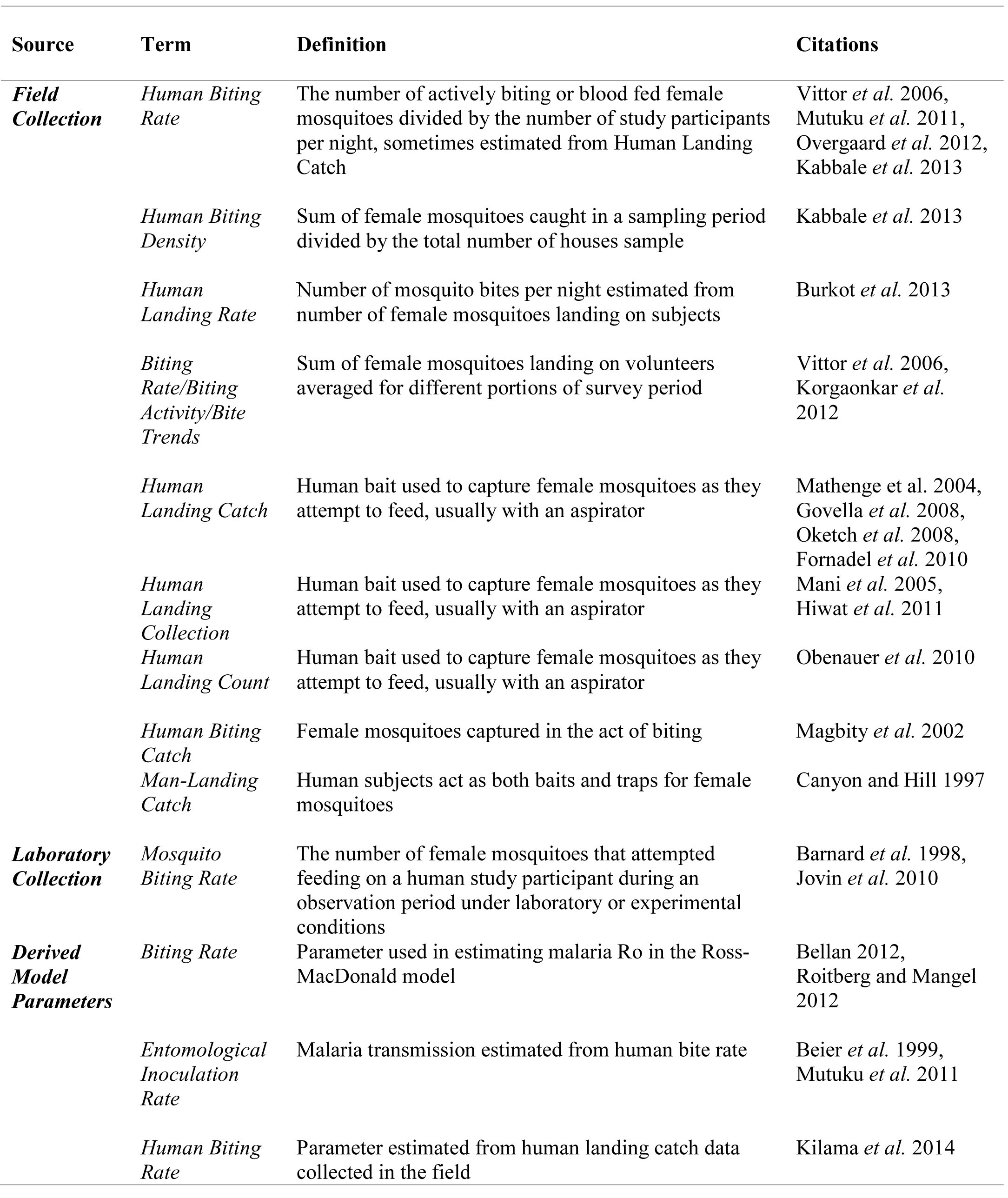
Glossary of terms related to mosquito biting activity used in the literature.

Ecuador’s southern El Oro province (Fig. 1) has been free of locally acquired malaria infections since 2011, although the mosquito species capable of vectoring *P. vivax* and *P. falciparum* malaria are still prevalent in the area [13]. Disease surveillance and control programs in developing countries typically suffer from limited resources in the face of high disease burden, however the Ecuadorian government has devoted a great deal of funding and logistic support to their Ministry of Health specifically for the detection and control of malaria following a resurgence of the disease in the late 1990’s, which has been previously described in detail [13]. Nevertheless, with recent outbreaks of malaria occurring in other Ecuadorian provinces and neighboring countries, the potential for re-emergence of malaria in El Oro creates a need to estimate the potential for malaria transmission as part of a surveillance system, and the behavior of blood-seeking female mosquitoes recorded via HLC can enhance the understanding of outbreak and exposure risks by illuminating relevant aspects of vector biology such as seasonal activity trends by species, peak biting activity by species, detailed shifts in species composition, and host seeking behavior and the propensity for endophagy (indoor feeding) [29–33]. This is information that can be directly incorporated into mosquito abatement strategies, surveillance protocols, and public education campaigns.

**Fig. 1.**
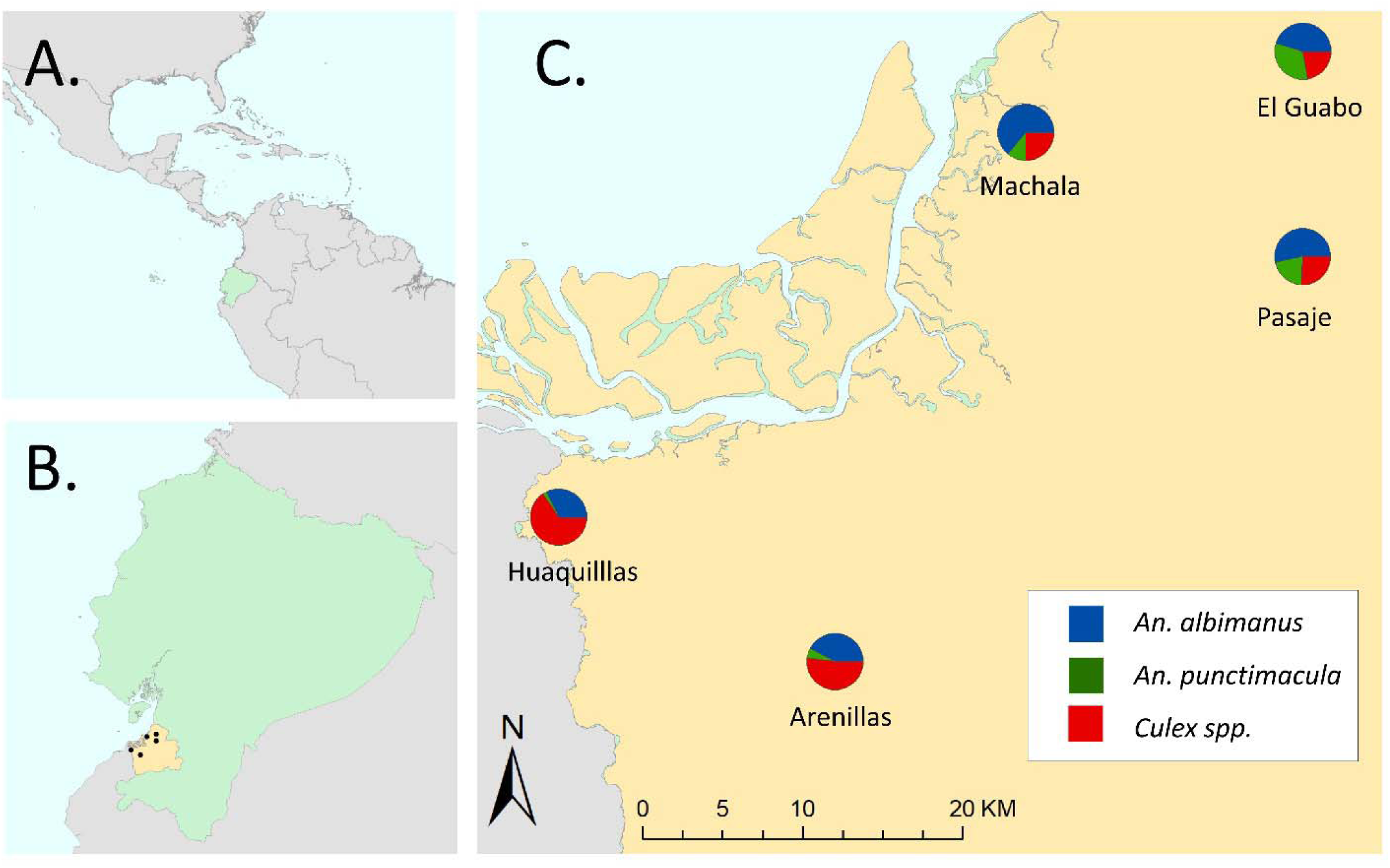
Data on mosquito biting rates were collected in five cities located in Ecuador’s (A) southern coastal El Oro province (B). Although the proportion of bites recorded relative to sampling effort for *Anopheles albimanus*, *An. Punctimacula*, and *Culex spp.* varied between cities, all three taxa of interest were detected across study sites (C).

Previous bite rate studies on *Anopheles* have demonstrated that mosquitoes can shift species composition and peak daily biting activity in response to abatement strategies, information that is crucial to developing and reviewing successful mosquito control efforts [21,34–36]. In Ecuador, there have been documented instances of epidemiological shifts in human disease patterns with concurrent transitions in species prevalence, and long-term collection of bite rate data at fine scales can capture these shifts [37]. This is an important consideration, as biting rate and peak biting activity are often considered as stable variables for any given species that can be directly reduced through routine interventions [18,24,38].

In this study, we examined nightly bite rate data collected in five cities from 2007-2012 in southern Ecuador. These data were collected as part of routine *Anopheline* surveillance by the National Service for the Control of Diseases Transmitted by Arthropod Vectors (SNEM) of the Ministry of Health. The goals of this paper are to 1) test the hypothesis that the bite indices for notable mosquito vectors in southern coastal Ecuador differ significantly across taxa 2) use an exploratory modeling framework to describe seasonal and diel variation in biting activity within each taxon and 3) use fine-scale spatial data to compare exophagic and endophagic feeding behaviors between taxa.

## Methods

### Bite Rate Data

Human landing catch (HLC) data were collected as a proxy for the biting activity (i.e. bite rate) of two malaria vectors (*Anopheles albimanus* and *Anopheles punctimacula*) and a pooled taxonomic grouping of potential arbovirus vectors (*Culex spp.*) at the household level from 2007 – 2012 in five coastal cities in Ecuador’s El Oro province: Huaquillas, Machala, El Guabo, Arenillas, and Pasaje (Fig. 1). In the first year of study, three primary sites (Huaquillas, Machala, and El Guabo) were surveyed every month to establish baseline data. In subsequent years, each site was surveyed four times annually, twice in the rainy season (January – May) and twice in the dry season. Field technicians were equipped with black stockings that covered the legs from the feet to above the knees and captured mosquitoes landing on the stockings with a mouth aspirator. Hourly collections were made each night (18:00 – 06:00) at study households, both inside homes and outdoors, allotting 50 minutes of each hour for aspiration and 10 minutes for specimen processing. All mosquitoes collected were brought back to the laboratory for counting, sexing, and species identification. Although sampling effort (i.e. number of survey nights) varied between cities (Arenillas (n=17), El Guabo (n=27), Huaquillas (n=38), Machala (n=33), Pasaje (n=2)), all three mosquito taxa were detected in all study sites (Fig. 1).

### Statistical Analysis

Regression models were used to determine if bite rates were fundamentally different for the three mosquito taxa, and to explore the influence of biting location (i.e. indoors vs. outdoors), season, and time of biting activity (i.e. hour of the day). Due to the size of the data set, we pooled across the 5 cities in the study. The bite rate data exhibited more zero observations than accommodated by commonly used error distributions for count data (e.g. Poisson or negative Binomial), an issue frequently encountered when modeling mosquito surveillance datasets, but not always treated in a statistically appropriate manner. We therefore used hurdle models, which combine a logistic regression model, the so-called hurdle, which describes the probability of being bitten at all, with a count model, which describes the number of bites conditional on being bitten [39]. In addition to wishing to use the appropriate statistics for the zero observations, we also use hurdle models rather than zero-inflated Poisson (ZIP) models, because we cannot distinguish between "structural" and "sampling" zeroes in these data. In our specific case, this leads to superior interpretability, because we can directly model the probability of being bitten by a particular species.

Hurdle models were fitted using the package ‘pscl’ in R ver. 3.3.1 (R Core Team, 2016), specifying a negative binomial error distribution and a log link for the count component, and a binomial error distribution and a logit link for the hurdle [40]. Variable selection for hurdle models was conducted based on Akaike’s Information Criterion [41]. Confidence intervals for model predictions were obtained using non-parametric bootstrapping with the ‘boot’ package in R [42,43].

## Results

We found that biting behavior for *An. albimanus, An. punctimacula,* and *Culex spp.* differed, both in terms of whether or not bites occurred (i.e. the odds (OR) of being bitten) and the number of bites per hour conditional on being bitten (expressed as incidence rate ratios, RR; Table 2). *An. albimanus* was the species most commonly observed biting (Fig. 3). The occurrence of *An. albimanus* bites in a given hour was four times as likely as no bites (OR 4.04, p< 0.001), with an average of 4.7 bites per hour (RR 4.74, p < 0.001).

**Table 2.** Species and location effects of a hurdle model of hourly biting rates. Model coefficients are presented as incidence rate ratios for the count model (which models hourly bites conditional on being bitten), and as odds ratios for the zero model (which models the probability of being bitten). A full table including the species-specific temporally resolved model coefficients is presented in the supplementary materials. Coefficients in this are representative for January at 6pm local time and relative *to An. albimanus* bite rates. Values in parentheses are 95% confidence intervals. Significance levels are P < 0.05 *, P < 0.01 **, P < 0.001 ***.

|  | Count model rate ratio | Zero model odds ratio |
| --- | --- | --- |
| Intercept (*An. albimanus*) | 4.74 (3.05-7.36) *** | 4.04 (2.39-6.82) *** |
| *Culex* spp. | 1.38 (0.79-2.42) | 3.31 (1.58-6.92) ** |
| *An. punctimacula* | 0.6 (0.31-1.18) | 0.65 (0.31-1.36) |
| Location outdoors | 1.55 (1.36-1.75) *** | 2.32 (2.03-2.64) *** |
| *Culex* : outdoors | 0.79 (0.66-0.94) ** | 0.58 (0.48-0.7) *** |
| *An. puncti.* : outdoors | 0.9 (0.71-1.13) | 0.8 (0.66-0.97) * |

Being outdoors more than doubled the odds of being bitten by *An. albimanus* (OR 2.32, p < 0.001), and increased the number of bites received when bitten by about 50% (RR 1.55, p < 0.001). For *Culex spp.* the odds of being bitten were lower overall (Fig. 3), albeit higher at the temporal reference levels of the model (i.e. January at 6pm) with an odds ratio of being bitten by *Culex* of 13.27 (p < 0.01) and an average of 6.5 bites when bitten (n.s. compared to *An. albimanus*). Being outdoors increased the odds of being bitten by *Culex* by about a third (OR 1.35, p < 0.01), and number of bites received by about a quarter (RR 1.22, P < 0.01), both to a lower extent than the associated increases for *An. albimanus*.

Bite rates for *An. punctimacula* were the lowest overall (Fig. 3), with a baseline odds ratio of being bitten of 2.62 and 2.94 bites per hour, but these base rates did not differ significantly from those for *An. albimanus*. Being outdoors increased the risk of being bitten by *An. albimanus* by about 80% (or 1.86, P < 0.05), and receiving bites by 40% (RR 1.40, n.s. compared to *An. albimanus*).

Months of peak high and low biting activity varied for the three taxa; the highest and lowest respective months for significant biting activity were March and July for *An. albimanus,* July and August for *An. punctimacula*, and February and July for *Culex spp* (Table 3).

## Discussion

Using data collected during a five year period across five cities in southern Ecuador, we found that there are spatial and temporal differences in the biting activity of mosquito taxa, including two species of known medical significance in Ecuador [13,44,45]. *An. albimanus*, a noted vector of malaria in Latin America, was the species most frequently observed attempting to bite human subjects, and although the baseline odds of being bitten by this species did not differ significantly from the other malaria vector, *An. punctimacula*, there are still distinct patterns of seasonal and temporal biting activity between the species (Table 2 & 3, S1). Despite these observed differences, all taxa demonstrated exophagic feeding tendencies, as being outside of households increased the risk of exposure to mosquito bites regardless of species (Table 3).

**Table 3.** Predicted average nightly bite rates (bites/hour) and associated 95% confidence intervals

| Month | *An. albimanus* |  | *An. punctimacula* |  | Culex spp. |  |
| --- | --- | --- | --- | --- | --- | --- |
|  | Indoors | Outdoors | Indoors | Outdoors | Indoors | Outdoors |
| Jan | 4.85<br>(2.57-8.01) | 8.06<br>(4.98-12.52) | 3.59<br>(1.53-6.06) | 5.26<br>(2.51-8.69) | 9.28<br>(5.85-11.99) | 11.31<br>(7.34-14.64) |
| Feb | 1.93<br>(0.78-3.58) | 3.73<br>(1.84-6.32) | 0.68<br>(0.19-1.27) | 1.29<br>(0.38-2.37) | 8.12<br>(5.28-10.43) | 9.8<br>(6.51-12.63) |
| Mar | 9.73<br>(5.42-15.88) | 15.96<br>(10.05-24.75) | 1.03<br>(0.34-1.77) | 1.69<br>(0.64-2.85) | 2.69<br>(1.33-3.8) | 3.45<br>(1.8-4.69) |
| Apr | 3.3<br>(1.66-5.54) | 5.6<br>(3.34-8.78) | 0.38<br>(0.11-0.68) | 0.71<br>(0.21-1.28) | 3.89<br>(2.24-5.01) | 4.81<br>(2.9-6.13) |
| May | 3.13<br>(1.6-5.21) | 5.26<br>(3.18-8.19) | 0.07<br>(0.03-0.11) | 0.13<br>(0.05-0.18) | 0.93<br>(0.39-1.49) | 1.23<br>(0.55-1.9) |
| Jun | 5.31<br>(3.18-8.29) | 8.31<br>(5.5-12.48) | 0.24<br>(0.08-0.4) | 0.43<br>(0.14-0.72) | 4.4<br>(2.93-5.54) | 5.25<br>(3.58-6.63) |
| Jul | 2.85<br>(1.49-4.67) | 4.71<br>(2.9-7.24) | 1.69<br>(0.78-2.67) | 2.39<br>(1.24-3.72) | 0.77<br>(0.35-1.18) | 0.99<br>(0.47-1.46) |
| Aug | 1.69<br>(0.65-3.22) | 3.38<br>(1.59-5.88) | 0.58<br>(0.17-1.05) | 1.06<br>(0.33-1.9) | 2.47<br>(1.19-3.53) | 3.18<br>(1.63-4.38) |
| Sep | 7.27<br>(3.56-12.53) | 12.68<br>(7.41-20.35) | 1.87<br>(0.53-3.52) | 3.34<br>(1.08-6.11) | 3.35<br>(1.6-4.8) | 4.35<br>(2.2-5.99) |
| Oct | 6.59<br>(3.44-11.02) | 11.1<br>(6.77-17.44) | 2.98<br>(0.71-6.01) | 5.88<br>(1.56-11.54) | 1.24<br>(0.57-1.87) | 1.6<br>(0.78-2.33) |
| Nov | 3.14<br>(1.38-5.63) | 5.77<br>(3.09-9.51) | 1.32<br>(0.33-2.59) | 2.58<br>(0.71-4.95) | 3.32<br>(1.7-4.57) | 4.23<br>(2.28-5.61) |
| Dec | 1.9<br>(0.95-3.15) | 3.19<br>(1.9-4.93) | 1.06<br>(0.37-1.79) | 1.69<br>(0.68-2.79) | 2.16<br>(0.83-3.6) | 2.99<br>(1.21-4.8) |

These findings have clear implications for the delivery of mosquito abatement services and the development of public outreach programs, as risk of exposure to mosquito bites is a demonstrated function of time (e.g. month, hour of activity), location (i.e. indoors vs. outdoors), and species of vector (Figs. 2 & 3). The hot rainy season occurs from January to April, and historically, malaria season was around March – July, peaking in May [13]. Given that we saw highest biting activity for *An. albimanus* in March, and lowest in July, but highest in July and lowest in August for *An. punctimacula*, the human exposure to these anopheline biting habits suggests a mix of activity level between the two species during the malaria season. For areas such as El Oro province, where malaria has been eliminated, *a priori* knowledge of exposure risks can be incorporated into a framework of targeted surveillance and control to prevent reemergence or reestablishment of malaria in the region. There is active vector control (household spraying) year round in Ecuador, but mosquito control efforts intensify immediately before and during the rainy season (January – May), when increased water availability provides ample habitat for the aquatic larval stages of mosquitoes. However, such interventions are either focused on reducing overall mosquito abundance or targeted on pooled taxonomic groupings (e.g. managing malarial infections by treating the genus *Anopheles* as a single group). We found that the biting activity of the primary malaria vectors extends beyond the spray season – particularly *An. puntimacula*, which has peak activity a full 2 months after spraying is finished. This could potentially allow additional malaria activity later in the season, and increase the role of the vector thought to be less important in Latin America. Incorporation of temporal biting trends by species into management plans (i.e. peak months of biting activity) has the potential to increase the effectiveness and efficiency of mosquito control programs by allowing decision makers to focus resources at time periods critical to disrupting life cycles of particular vectors, and consequently the diseases they spread.

**Fig. 2.**
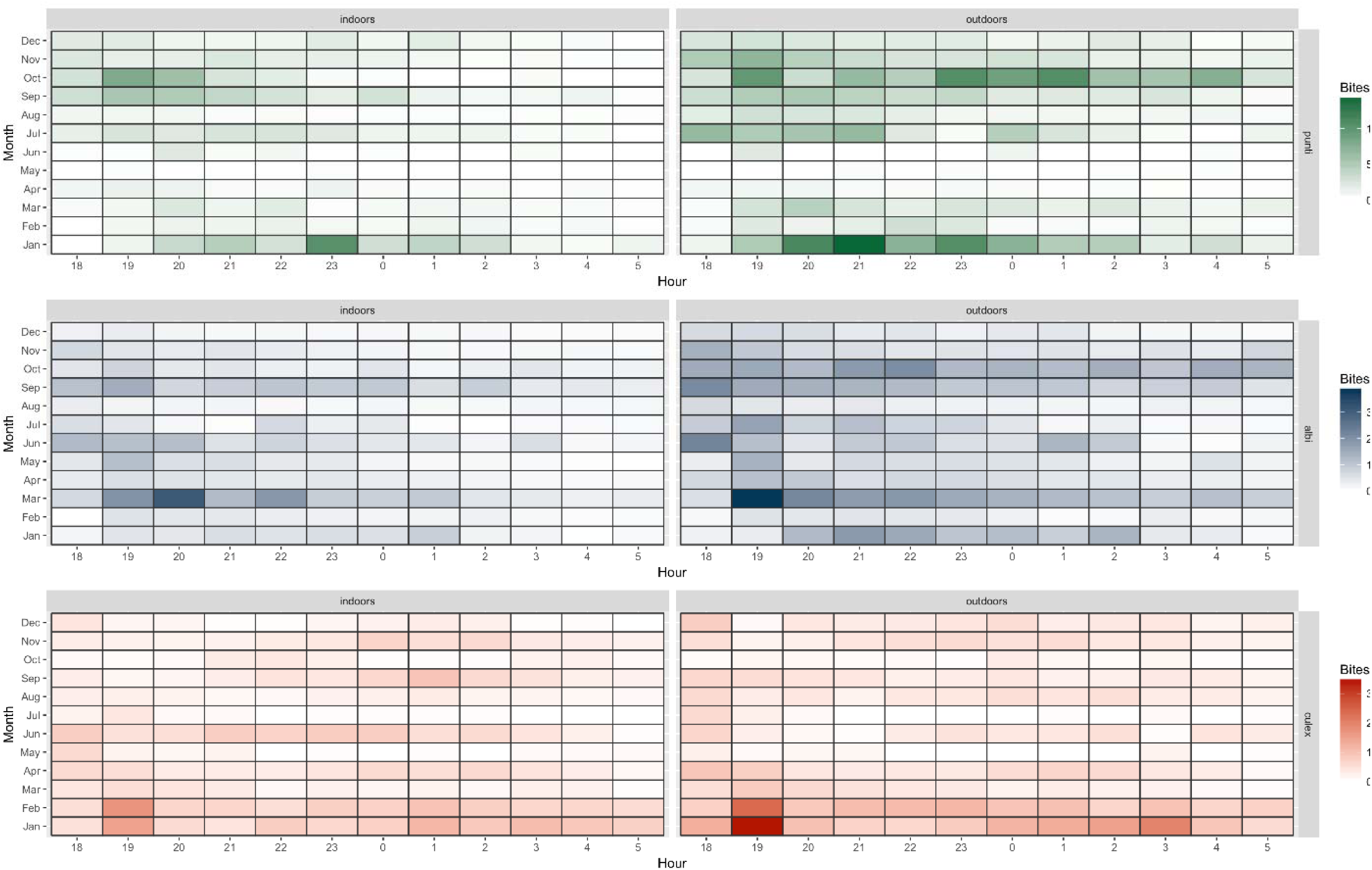
Raw observations of average hourly bite rates by species and location.

**Fig. 3.**
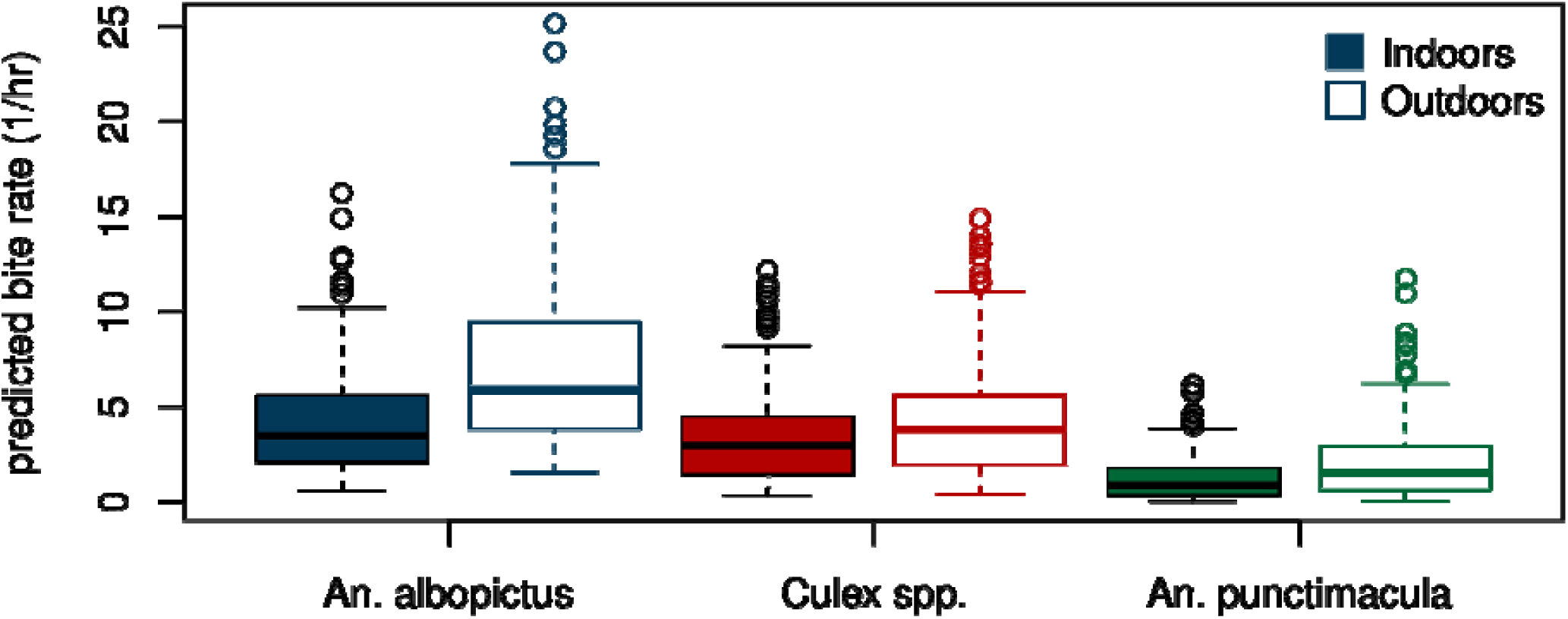
Hourly bite rates by species and location as predicted by the hurdle model across all months and hours of the night.

The dynamics of malaria transmission in Latin American countries are complex, and to fully understand localized disease risks we ultimately need to examine not only exposure to vectors but also the vectorial capacity of mosquitoes, which can vary with species and environment [46– 48]. That said, quantifying taxonomic-specific biting patterns is still a useful endeavor when developing control strategies, as demonstrably competent disease vectors are known to display differential feeding behaviors throughout their geographic ranges. This is the case with *An. albimanus*, which has been observed displaying both anthropophilic and zoophilic feeding preferences depending on location, potentially responsible for spatial variability in the true risk of disease transmission to humans [2,49–51]. Similarly, patterns of microhabitat use can vary spatially, with the proportion of endophagic versus exophagic mosquitoes depending not only on taxon, but also spatially contextual factors such as environment and housing structures [32,51]. In these instances, the collection of HLC data can serve as a better indicator of true exposure risk than simply documenting the presence of known competent vectors.

The utility of bite rate indices as a relatively low-cost surveillance tool is well documented [22,23,50]. However, the ability to differentiate closely related mosquito species may serve as an additional logistical challenge to the field surveillance of mosquito vectors in Ecuador. Female *An. punctimacula* are morphologically similar to *An. calderoni*, another vector of malaria in Latin America [52]. Despite being a competent vector of *Plasmodium spp.*, *An. calderoni* was only recently confirmed in several Latin American countries, including Ecuador, due to the systematic misclassification of the species [52,53]. The potential for misidentification of these taxa on surveys may obscure true species-level patterns in biting activity.

The bite count data in this study were collected at a very high temporal (e.g. hourly) and spatial (e.g. inside and outside of households) resolutions but were pooled across the five study cities for statistical analysis. This was largely due to the high number of variable combinations (e.g. species by month, species by hour) relative to the number of collection nights and the inherent zero-inflated nature of count data. Ideally, future studies would strive for more spatio-temporally balanced data collection across cities, allowing for more robust exploration of the spatial variation in biting trends across the study region. This would involve deploying multiple trained teams, which may be a prohibitive constraint at present. Despite these limitations, human bite rate indices remain a valuable tool in the collection of high-resolution vector ecology data, enabling quantification of risks associated with exposure to mosquito bites in a manner that is cost-effective and simple to implement.

## Conclusions

This is the first time that fine-scale spatial and temporal differences in the biting patterns of mosquito taxa have been reported for El Oro province in southern coastal Ecuador. These findings provide detailed information for targeting vector control and household level behavioral interventions. The data used to examine human biting trends were collected as part of routine vector surveillance conducted by the Ministry of Health, but such data have not been collected since the end of this dataset. As we have learned from experiences with dengue in the region, even when there is decline in cases, as happened prior to the 1970s, relaxing vector control, and reducing surveillance, can lead to rapid reemergence. Reinstating such surveillance measures will provide important information that will aid in preventing malaria re-emergence.

## List of Abbreviations

HLC: human landing catch
EIR: entomological inoculation rate
HBR: human biting rate
LR: landing rate
SNEM: National Service for the Control of Diseases Transmitted by Arthropod Vectors (Ecuador)
OR: odds ratio
RR: rate ratio

## Declarations

### Ethics approval and consent to participate

Not applicable

### Consent for publication

Not applicable

### Availability of data and material

The original datasets analysed during the current study are not publicly available, as they belong to SNEM - National Service for the Control of Diseases Transmitted by Arthropod Vectors, and the Ministry of Health, of Ecuador. Derived datasets are available on request from the corresponding author.

### Authors’ contributions

AMS, MS, EBA conceived the project and designed the surveys. SJR, CAL, PHBS conducted statistical analyses and prepared figures and maps. SJR, CAL, PHBS drafted the manuscript. LK, JTK, JA, MS contributed to the development of the database. SJR, CAL, PHBS, NH, MS, JA, LFN, EBA, ME, DA, JTK, LK, LF, and AMS contributed to the final version of the manuscript. All authors read and approved the final manuscript.

## Acknowledgments

We thank the Ministry of Health, SNEM, and the vector teams who contributed to this study. We thank the people of Huquillas, Arenilla, El Guabo, Pasaje, and Machala for participating in the original surveillance work.

## Supplementary Materials

**Supplementary Table 1:** Hurdle model of hourly biting rates. Model coefficients are presented as incidence rate ratios for the count model (which models hourly bites conditional on being bitten using a negative binomial error distribution and log link), and as odds ratios for the zero model (which models the probability of being bitten using binomial errors and a logit link). Values in parentheses are 95% confidence intervals. Significance levels are P < 0.05 *, P < 0.01 **, P < 0.001 ***.

|  | Count model rate ratio | Zero model odds ratio |
| --- | --- | --- |
| (Intercept) | 4.74 (3.05-7.36) *** | 4.04 (2.39-6.82) *** |
| Speciesculex | 1.38 (0.79-2.42) | 3.31 (1.58-6.92) ** |
| Speciespunti | 0.6 (0.31-1.18) | 0.65 (0.31-1.36) |
| Locationoutdoors | 1.55 (1.36-1.75) *** | 2.32 (2.03-2.64) *** |
| MonthFeb | 0.52 (0.34-0.79) ** | 0.31 (0.19-0.52) *** |
| MonthMar | 2.18 (1.4-3.39) *** | 1.4 (0.79-2.49) |
| MonthApr | 0.66 (0.45-0.99) * | 0.74 (0.45-1.22) |
| MonthMay | 0.6 (0.36-1) . | 0.78 (0.42-1.46) |
| MonthJun | 0.96 (0.56-1.65) | 1.73 (0.81-3.68) |
| MonthJul | 0.51 (0.26-1) * | 0.83 (0.37-1.88) |
| MonthAug | 0.5 (0.33-0.75) *** | 0.26 (0.16-0.42) *** |
| MonthSep | 1.82 (1.22-2.73) ** | 0.8 (0.48-1.32) |
| MonthOct | 1.49 (0.87-2.55) | 1 (0.51-1.97) |
| MonthNov | 0.79 (0.53-1.17) | 0.47 (0.29-0.76) ** |
| MonthDec | 0.32 (0.21-0.48) *** | 0.62 (0.37-1.03) . |
| Hour_factor19 | 1.05 (0.8-1.37) | 1.42 (1-2.02) * |
| Hour_factor20 | 0.8 (0.61-1.06) | 1.17 (0.83-1.64) |
| Hour_factor21 | 0.77 (0.58-1.01) . | 0.84 (0.61-1.18) |
| Hour_factor22 | 0.7 (0.53-0.93) * | 0.91 (0.65-1.28) |
| Hour_factor23 | 0.66 (0.5-0.89) ** | 0.59 (0.43-0.82) ** |
| Hour_factor0 | 0.6 (0.45-0.8) *** | 0.6 (0.44-0.83) ** |
| Hour_factor1 | 0.59 (0.44-0.8) *** | 0.41 (0.3-0.57) *** |
| Hour_factor2 | 0.49 (0.36-0.66) *** | 0.39 (0.28-0.53) *** |
| Hour_factor3 | 0.45 (0.33-0.61) *** | 0.36 (0.26-0.49) *** |
| Hour_factor4 | 0.44 (0.32-0.6) *** | 0.27 (0.2-0.37) *** |
| Hour_factor5 | 0.48 (0.35-0.66) *** | 0.23 (0.16-0.31) *** |
| Speciesculex:Locationoutdoors | 0.79 (0.66-0.94) ** | 0.58 (0.48-0.7) *** |
| Speciespunti:Locationoutdoors | 0.9 (0.71-1.13) | 0.8 (0.66-0.97) * |
| Speciesculex:MonthFeb | 1.58 (0.92-2.73) . | 3.89 (1.87-8.12) *** |
| Speciespunti:MonthFeb | 0.88 (0.45-1.71) | 0.37 (0.18-0.76) ** |
| Speciesculex:MonthMar | 0.15 (0.08-0.27) *** | 0.16 (0.08-0.35) *** |
| Speciespunti:MonthMar | 0.15 (0.08-0.3) *** | 0.2 (0.09-0.44) *** |
| Speciesculex:MonthApr | 0.58 (0.35-0.97) * | 0.65 (0.33-1.29) |
| Speciespunti:MonthApr | 0.35 (0.18-0.65) *** | 0.12 (0.06-0.24) *** |
| Speciesculex:MonthMay | 0.22 (0.08-0.58) ** | 0.11 (0.04-0.3) *** |
| Speciespunti:MonthMay | 0 (0-Inf) | 0.04 (0.01-0.13) *** |
| Speciesculex:MonthJun | 0.38 (0.19-0.79) ** | 0.65 (0.22-1.94) |
| Speciespunti:MonthJun | 0.09 (0.03-0.29) *** | 0.05 (0.02-0.13) *** |
| Speciesculex:MonthJul | 0.14 (0.05-0.41) *** | 0.13 (0.04-0.39) *** |
| Speciespunti:MonthJul | 0.64 (0.24-1.66) | 1.06 (0.34-3.31) |
| Speciesculex:MonthAug | 0.62 (0.37-1.04) . | 0.81 (0.41-1.6) |
| Speciespunti:MonthAug | 0.63 (0.34-1.16) | 0.48 (0.24-0.97) * |
| Speciesculex:MonthSep | 0.26 (0.15-0.43) *** | 0.26 (0.13-0.53) *** |
| Speciespunti:MonthSep | 0.57 (0.31-1.04) . | 0.28 (0.14-0.57) *** |
| Speciesculex:MonthOct | 0.1 (0.04-0.21) *** | 0.13 (0.05-0.33) *** |
| Speciespunti:MonthOct | 2.01 (0.79-5.1) | 0.13 (0.05-0.34) *** |
| Speciesculex:MonthNov | 0.5 (0.3-0.83) ** | 0.6 (0.3-1.17) |
| Speciespunti:MonthNov | 1.47 (0.81-2.68) | 0.25 (0.13-0.51) *** |
| Speciesculex:MonthDec | 1.67 (0.92-3.02) . | 0.13 (0.06-0.26) *** |
| Speciespunti:MonthDec | 0.92 (0.49-1.72) | 0.53 (0.26-1.1) . |
| Speciesculex:Hour_factor19 | 0.99 (0.67-1.48) | 0.49 (0.29-0.82) ** |
| Speciespunti:Hour_factor19 | 1.28 (0.79-2.08) | 0.94 (0.59-1.51) |
| Speciesculex:Hour_factor20 | 0.8 (0.53-1.19) | 0.49 (0.29-0.8) ** |
| Speciespunti:Hour_factor20 | 1.51 (0.93-2.46) . | 1.2 (0.75-1.91) |
| Speciesculex:Hour_factor21 | 0.94 (0.62-1.43) | 0.45 (0.27-0.73) ** |
| Speciespunti:Hour_factor21 | 1.24 (0.75-2.03) | 1.48 (0.94-2.35) . |
| Speciesculex:Hour_factor22 | 1.05 (0.7-1.58) | 0.49 (0.3-0.8) ** |
| Speciespunti:Hour_factor22 | 1.1 (0.67-1.8) | 1.33 (0.84-2.11) |
| Speciesculex:Hour_factor23 | 1.27 (0.84-1.92) | 0.86 (0.53-1.4) |
| Speciespunti:Hour_factor23 | 1.34 (0.8-2.24) | 1.65 (1.05-2.61) * |
| Speciesculex:Hour_factor0 | 1.86 (1.23-2.82) ** | 0.85 (0.52-1.38) |
| Speciespunti:Hour_factor0 | 0.89 (0.53-1.48) | 1.54 (0.98-2.43) . |
| Speciesculex:Hour_factor1 | 2.09 (1.37-3.2) *** | 1.03 (0.63-1.66) |
| Speciespunti:Hour_factor1 | 0.99 (0.58-1.7) | 1.76 (1.11-2.79) * |
| Speciesculex:Hour_factor2 | 2.24 (1.46-3.44) *** | 0.98 (0.61-1.59) |
| Speciespunti:Hour_factor2 | 0.98 (0.57-1.7) | 1.8 (1.14-2.85) * |
| Speciesculex:Hour_factor3 | 2.12 (1.37-3.27) *** | 0.87 (0.54-1.41) |
| Speciespunti:Hour_factor3 | 0.88 (0.5-1.54) | 1.81 (1.14-2.88) * |
| Speciesculex:Hour_factor4 | 1.71 (1.09-2.68) * | 0.84 (0.52-1.35) |
| Speciespunti:Hour_factor4 | 1.17 (0.63-2.15) | 1.47 (0.9-2.37) |
| Speciesculex:Hour_factor5 | 1.2 (0.75-1.92) | 0.72 (0.44-1.16) |
| Speciespunti:Hour_factor5 | 0.81 (0.41-1.59) | 1.13 (0.68-1.88) |

## References

1. Pan American Health Organization. Report on the situation of malaria in the Americas, 2014. Washington DC, US: PAHO; 2016 p. 114.

2. Sinka ME, Rubio-Palis Y, Manguin S, Patil AP, Temperley WH, Gething PW, et al. The dominant Anopheles vectors of human malaria in the Americas: occurrence data, distribution maps and bionomic précis. Parasit. Vectors. 2010;3:72.

3. Cruz LR, Spangenberg T, Lacerda MV, Wells TN. Malaria in South America: a drug discovery perspective. Malar. J. 2013;12:168.

4. CDC. Health Information for Travelers to Ecuador, Including the Galapages Islands [Internet]. Centers for Disease Control; 2016 [cited 2016 Dec 1]. Available from: http://wwwnc.cdc.gov/travel/destinations/traveler/none/ecuador

5. World Health Organization, World Health Organization, Global Malaria Programme. Disease surveillance for malaria control: an operational manual. [Internet]. Geneva, Switzerland: World Health Organization; 2012 [cited 2017 Jun 19]. Available from: http://apps.who.int/iris/bitstream/10665/44851/1/9789241503341_eng.pdf

6. Guedes DRD, Cordeiro MT, Melo-Santos M a. V, Magalhaes T, Marques E, Regis L, et al. Patient-based dengue virus surveillance in Aedes aegypti from Recife, Brazil. J. Vector Borne Dis. 2010;47:67–75.

7. Flies EJ, Toi C, Weinstein P, Doggett SL, Williams CR. Converting Mosquito Surveillance to Arbovirus Surveillance with Honey-Baited Nucleic Acid Preservation Cards. Vector-Borne Zoonotic Dis. 2015;15:397–403.

8. Lopez-Cevallos D, Chi C. Inequity in health care utilization in Ecuador: an analysis of current issues and potential solutions. Int. J. Equity Health. 2012;11:A6.

9. Stewart-Ibarra AM, Muñoz ÁG, Ryan SJ, Ayala EB, Borbor-Cordova MJ, Finkelstein JL, et al. Spatiotemporal clustering, climate periodicity, and social-ecological risk factors for dengue during an outbreak in Machala, Ecuador, in 2010. BMC Infect. Dis. [Internet]. 2014 [cited 2017 Apr 26];14. Available from: http://bmcinfectdis.biomedcentral.com/articles/10.1186/s12879-014-0610-4

10. Dumonteil E, Herrera C, Martini L, Grijalva MJ, Guevara AG, Costales JA, et al. Chagas Disease Has Not Been Controlled in Ecuador. Tanowitz HB, editor. PloS ONE. 2016;11:e0158145.

11. Stewart Ibarra AM, Ryan SJ, Beltrán E, Mejía R, Silva M, Muñoz Á. Dengue Vector Dynamics (Aedes aegypti) Influenced by Climate and Social Factors in Ecuador: Implications for Targeted Control. Mores CN, editor. PLoS ONE. 2013;8:e78263.

12. Sturrock HJW, Hsiang MS, Cohen JM, Smith DL, Greenhouse B, Bousema T, et al. Targeting Asymptomatic Malaria Infections: Active Surveillance in Control and Elimination. PLoS Med. 2013;10:e1001467.

13. Krisher LK, Krisher J, Ambuludi M, Arichabala A, Beltr？n-Ayala E, Navarrete P, et al. Successful malaria elimination in the Ecuador?Peru border region: epidemiology and lessons learned. Malar. J. [Internet]. 2016 [cited 2017 Jun 16];15. Available from: http://malariajournal.biomedcentral.com/articles/10.1186/s12936-016-1630-x

14. Mordecai EA, Paaijmans KP, Johnson LR, Balzer C, Ben-Horin T, de Moor E, et al. Optimal temperature for malaria transmission is dramatically lower than previously predicted. Thrall P, editor. Ecol. Lett. 2013;16:22–30.

15. Ross R. Some a priori pathometric equations. Br. Med. J. 1915;1:546–7.

16. MacDonald G. The Epidemiology and Control of Malaria. 1957;xv + 201 + xl + 11 pp.

17. Mordecai EA, Cohen JM, Evans MV, Gudapati P, Johnson LR, Lippi CA, et al. Detecting the impact of temperature on transmission of Zika, dengue, and chikungunya using mechanistic models. Althouse B, editor. PLoS Negl. Trop. Dis. 2017;11:e0005568.

18. Smith DL, McKenzie FE. Statics and dynamics of malaria infection in Anopheles mosquitoes. Malar. J. 2004;3:13.

19. Kelly-Hope LA, McKenzie FE. The multiplicity of malaria transmission: a review of entomological inoculation rate measurements and methods across sub-Saharan Africa. Malar. J. 2009;8:19.

20. Kang S-H. Comparative Repellency of Essential Oils against Culex pipiens pallens (Diptera: Culicidae). J. Korean Soc. Appl. Biol. Chem. 2009;52:353–9.

21. Mutuku FM, King CH, Mungai P, Mbogo C, Mwangangi J, Muchiri EM, et al. Impact of insecticide-treated bed nets on malaria transmission indices on the south coast of Kenya. Malar. J. 2011;10:356.

22. Overgaard HJ, Sæbø S, Reddy MR, Reddy VP, Abaga S, Matias A, et al. Light traps fail to estimate reliable malaria mosquito biting rates on Bioko Island, Equatorial Guinea. Malar. J. 2012;11:56.

23. Kumar A, Kabadi D, Korgaonkar N, Yadav R, Dash A. Mosquito biting activity on humans & detection of Plasmodium falciparum infection in Anopheles stephensi in Goa, India. Indian J. Med. Res. 2012;135:120.

24. Bellan SE. The Importance of Age Dependent Mortality and the Extrinsic Incubation Period in Models of Mosquito-Borne Disease Transmission and Control. Cornell SJ, editor. PLoS ONE. 2010;5:e10165.

25. Onyango SA, Kitron U, Mungai P, Muchiri EM, Kokwaro E, King CH, et al. Monitoring malaria vector control interventions: effectiveness of five different adult mosquito sampling methods. J. Med. Entomol. 2013;50:1140–51.

26. World Health Organization. Malaria entomology and vector control guide for participants. Geneva, Switzerland: World Health Organization; 2013.

27. Sikaala CH, Killeen GF, Chanda J, Chinula D, Miller JM, Russell TL, et al. Evaluation of alternative mosquito sampling methods for malaria vectors in Lowland South - East Zambia. Parasit. Vectors. 2013;6:91.

28. Kenea O, Balkew M, Tekie H, Gebre-Michael T, Deressa W, Loha E, et al. Comparison of two adult mosquito sampling methods with human landing catches in south-central Ethiopia. Malar. J. [Internet]. 2017 [cited 2017 Jun 19];16. Available from: http://malariajournal.biomedcentral.com/articles/10.1186/s12936-016-1668-9

29. Herrera-Varela M, Orjuela LI, Pe?alver C, Conn JE, Qui?ones ML. Anopheles species composition explains differences in Plasmodium transmission in La Guajira, northern Colombia. Memrias Inst. Oswaldo Cruz. 2014;109:952–6.

30. Loaiza JR, Bermingham E, Scott ME, Rovira JR, Conn JE. Species composition and distribution of adult Anopheles (Diptera: Culicidae) in Panama. J. Med. Entomol. 2008;45:841–51.

31. Vittor AY, Gilman RH, Tielsch J, Glass GE, Shields T, Lozano WS, et al. The effect of deforestation on the human-biting rate of Anopheles darlingi, the primary vector of Falciparum malaria in the Peruvian Amazon. Am. J. Trop. Med. Hyg. 2006;74:3–11.

32. Hobbs JH, Sexton JD, St Jean Y, Jacques JR. The biting and resting behavior of Anopheles albimanus in northern Haiti. J. Am. Mosq. Control Assoc. 1986;2:150–3.

33. Sáenz FE, Morton LC, Okoth SA, Valenzuela G, Vera-Arias CA, Vélez-Álvarez E, et al. Clonal population expansion in an outbreak of Plasmodium falciparum on the northwest coast of Ecuador. Malar. J. [Internet]. 2015 [cited 2017 Jul 7];14. Available from: http://www.malariajournal.com/content/14/1/497

34. Russell TL, Govella NJ, Azizi S, Drakeley CJ, Kachur SP, Killeen GF. Increased proportions of outdoor feeding among residual malaria vector populations following increased use of insecticide-treated nets in rural Tanzania. Malar. J. 2011;10:80.

35. Sougoufara S, Di?dhiou S, Doucour? S, Diagne N, Semb?ne P, Harry M, et al. Biting by Anopheles funestus in broad daylight after use of long-lasting insecticidal nets: a new challenge to malaria elimination. Malar. J. 2014;13:125.

36. Lindblade KA, Gimnig JE, Kamau L, Hawley WA, Odhiambo F, Olang G, et al. Impact of Sustained Use of Insecticide-Treated Bednets on Malaria Vector Species Distribution and Culicine Mosquitoes. J. Med. Entomol. 2006;43:428–32.

37. Cifuentes SG, Trostle J, Trueba G, Milbrath M, Baldeón ME, Coloma J, et al. Transition in the Cause of Fever from Malaria to Dengue, Northwestern Ecuador, 1990–2011. Emerg. Infect. Dis. 2013;19:1642–5.

38. Kabbale FG, Akol AM, Kaddu JB, Onapa AW. Biting patterns and seasonality of anopheles gambiae sensu lato and anopheles funestus mosquitoes in Kamuli District, Uganda. Parasit. Vectors. 2013;6:340.

39. Zeileis A, Kleiber C, Jackman S. Regression Models for Count Data in *R*. J. Stat. Softw. [Internet]. 2008 [cited 2017 Jun 9];27. Available from: http://www.jstatsoft.org/v27/i08/

40. Jackman S. pscl: Classes and methods for R developed in the Political Science Computational Library, Stanford University. Stanford, California; 2015.

41. Burnham KP, Anderson DR, Burnham KP. Model selection and multimodel inference: a practical information-theoretic approach. 2nd ed. New York: Springer; 2002.

42. Davison AC, Hinkley DV. Bootstrap methods and their application. Cambridge; New York, NY, USA: Cambridge University Press; 1997.

43. Canty A, Ripley B. boot: Bootstratp R (S-Plus) Functions. 2017.

44. Pinault LL, Hunter FF. Characterization of larval habitats of Anopheles albimanus, Anopheles pseudopunctipennis, Anopheles punctimacula, and Anopheles oswaldoi s.l. populations in lowland and highland Ecuador. J. Vector Ecol. 2012;37:124–36.

45. Rubio-Palis Y, Zimmerman RH. Ecoregional classification of malaria vectors in the neotropics. J. Med. Entomol. 1997;34:499–510.

46. Turell MJ. Effect of environmental temperature on the vector competence of Aedes taeniorhynchus for Rift Valley fever and Venezuelan equine encephalitis viruses. Am. J. Trop. Med. Hyg. 1993;49:672–6.

47. Okech BA, Gouagna LC, Yan G, Githure JI, Beier JC. Larval habitats of Anopheles gambiae s.s. (Diptera: Culicidae) influences vector competence to Plasmodium falciparum parasites. Malar. J. 2007;6:50.

48. Antonio-nkondjio C, Kerah CH, Simard F, Awono-ambene P, Chouaibou M, Tchuinkam T, et al. Complexity of the Malaria Vectorial System in Cameroon: Contribution of Secondary Vectors to Malaria Transmission. J. Med. Entomol. 2006;43:1215–21.

49. Loyola EG, González-Cerón L, Rodríguez MH, Arredondo-Jiménez JI, Bennett S, Bown DN. Anopheles albimanus (Diptera: Culicidae) host selection patterns in three ecological areas of the coastal plains of Chiapas, southern Mexico. J. Med. Entomol. 1993;30:518–23.

50. Solarte Y, Hurtado C, Gonzalez R, Alexander B. Man-biting activity of Anopheles (Nyssorhynchus) albimanus and An. (Kerteszia) neivai (Diptera: Culicidae) in the Pacific lowlands of Colombia. Mem. Inst. Oswaldo Cruz. 1996;91:141–6.

51. Bown DN, Rodriguez MH, Arredondo-Jimenez JI, Loyola EG, Rodriguez MC. Intradomiciliary behavior of Anopheles albimanus on the coastal plain of southern Mexico: implications for malaria control. J. Am. Mosq. Control Assoc. 1993;9:321–4.

52. González R, Carrejo N, Wilkerson RC, Alarcon J, Alarcon-Ormasa J, Ruiz F, et al. Confirmation of Anopheles (Anopheles) calderoni Wilkerson, 1991 (Diptera: Culicidae) in Colombia and Ecuador through molecular and morphological correlation with topotypic material. Mem. Inst. Oswaldo Cruz. 2010;105:1001–9.

53. Orjuela LI, Ahumada ML, Avila I, Herrera S, Beier JC, Quiñones ML. Human biting activity, spatial–temporal distribution and malaria vector role of Anopheles calderoni in the southwest of Colombia. Malar. J. [Internet]. 2015 [cited 2017 Jun 16];14. Available from: http://www.malariajournal.com/content/14/1/256

